# The polyol pathway is an evolutionarily conserved system for sensing glucose uptake

**DOI:** 10.1101/2021.09.14.460366

**Authors:** Hiroko Sano, Akira Nakamura, Mariko Yamane, Hitoshi Niwa, Takashi Nishimura, Kimi Araki, Kazumasa Takemoto, Kei-ichiro Ishiguro, Hiroki Aoki, Masayasu Kojima

## Abstract

Cells must adjust the expression levels of metabolic enzymes in response to fluctuating nutrient supply. For glucose, such metabolic remodeling is highly dependent on a master transcription factor ChREBP/MondoA. However, it remains elusive how glucose fluctuations are sensed by ChREBP/MondoA despite the stability of major glycolytic pathways. Here we show that in both flies and mice, ChREBP/MondoA activation in response to glucose ingestion depends on an evolutionarily conserved glucose-metabolizing pathway: the polyol pathway. The polyol pathway converts glucose to fructose via sorbitol. It has been believed that this pathway is almost silent, and its activation in hyperglycemic conditions has deleterious effects on human health. We show that the polyol pathway is required for the glucose-induced nuclear translocation of Mondo, a *Drosophila* homologue of ChREBP/MondoA, which directs gene expression for organismal growth and metabolism. Likewise, inhibition of the polyol pathway in mice impairs ChREBP’s nuclear localization and reduces glucose tolerance. We propose that the polyol pathway is an evolutionarily conserved sensing system for the glucose uptake that allows metabolic remodeling.

## Introduction

The accurate sensing of ingested nutrients is vital for organismal survival. Animals need to sense quantitative and temporal changes in their nutritional status due to daily feeding to optimize metabolism. Glucose is the most commonly used energy source for animals and provides a good example of how they developed systems that achieve nutritional adaptation. Ingestion of glucose induces nutritional adaptation in the form of increases in glucose absorption and metabolism as well as lipogenesis to store excess nutrients, and inadequate adaptation might contribute to metabolic diseases such as obesity and type 2 diabetes. Most of glucose-induced nutritional adaptation is the results of glucose-dependent transcription (Towle, 2005), and such metabolic remodeling largely relies on a master transcription factor, carbohydrate responsive element binding protein (ChREBP) (Richards et al., 2017). ChREBP activates the expression of glycolytic and lipogenic genes with their obligated partner, Max-like protein X (Mlx), thereby storing excess nutrients in the form of lipids in the liver and adipose tissues (Iizuka et al., 2004). MondoA, a paralog of ChREBP, functions in the skeletal muscle (Billin et al., 2000). In *Drosophila*, the homologue of ChREBP/MondoA is encoded by a single gene, *Mondo*. Transcriptome analysis has shown that the Mondo-Mlx (also called Bigmax in *Drosophila*) complex induces global changes in metabolic gene expression according to sugar uptake (Havula et al., 2013; Mattila et al., 2015). To achieve a metabolic shift, information on glucose availability must somehow be transmitted to ChREBP/MondoA.

Upon glucose uptake, ChREBP/MondoA is translocated to the nucleus, which is a pivotal step for ChREBP/MondoA activation. ChREBP/MondoA shuttles between cytoplasmic and nuclear compartments and is primarily localized at the cytoplasm in the basal state (Billin et al., 2000; Davies et al., 2008; Kawaguchi et al., 2001). Glucose stimuli shift ChREBP/MondoA to the nuclei through the N-terminal domain containing a nuclear localization signal (Davies et al., 2010; Li et al., 2006). This process is critical to exert their transcriptional activities. Although the precise mechanism is unknown, it has been thought that metabolites generated from glucose directly or indirectly regulate the nuclear localization of ChREBP/MondoA.

Using nuclear translocation and transcriptional activation as indicators, ChREBP/MondoA-activating sugars have been explored by administering candidate sugars to cultured cells. So far, several candidates such as xylulose 5-phosphate (Xu5P), glucose-6-phosphate and fructose-2,6-bisphosphate have been identified (Arden et al., 2012; Dentin et al., 2012; Diaz-Moralli et al., 2012; Iizuka et al., 2013; Kabashima et al., 2003; Li et al., 2010; Peterson et al., 2010; Petrie et al., 2013; Stoltzman et al., 2008). These sugars are synthesized through either glycolysis or the pentose phosphate pathway (PPP) that branches off from glycolysis. Thus, cells were thought to detect blood glucose levels by assessing the activities of these two pathways. However, the levels of metabolites in these pathways remain mostly constant after glucose uptake partly due to the storage sugars (Peeters et al., 2017). Storage sugars such as glycogen are known to provide buffering action to prevent drastic changes in glucose metabolism; excessive nutrient uptake promotes the conversion of glucose-6-phosphate into glycogen, while starvation induces the breakdown of glycogen into glucose-6-phosphate. Moreover, glycolysis is tightly regulated by feedback control; glycolytic enzymes, including hexokinase working at the most upstream point in the pathway, are activated or inhibited by downstream metabolites (Berg, 2006). Therefore, glycolysis and PPP are likely to be inadequate as real-time sensors to detect small changes in glucose concentration under normal physiological conditions. These findings suggest that the activation of ChREBP/MondoA involves a hitherto unrecognized pathway.

In this study we show that the polyol pathway is required for the activation of *Drosophila* Mondo and mammalian ChREBP using genetic inhibition of this pathway. The polyol pathway is a two-step metabolic pathway, in which glucose is reduced to sorbitol then converted to fructose (Hers, 1956). It has long been believed that the polyol pathway is almost silent under normal physiological conditions but becomes active and harmful under hyperglycemic conditions (Brownlee, 2001; Lorenzi, 2007). However, the genes encoding polyol pathway enzymes are conserved from yeasts to humans, even though it is dispensable for the synthesis of ATP or biomolecules, suggesting that the polyol pathway plays an important previously unknown role across species. We demonstrate that the polyol pathway metabolites promote, and its mutation disturbs nuclear translocation of Mondo/ChREBP in *Drosophila* and mice. The polyol pathway is required for the regulation of Mondo/ChREBP-target metabolic genes, leading to proper growth and physiology. Our results show that the polyol pathway is an evolutionarily conserved system for sensing glucose uptake that allows metabolic remodeling.

## Results

### The polyol pathway is required for Mondo-mediated *CCHa2* expression

As a marker to assess what might activates Mondo/ChREBP, we chose a glucose-responsive hormone, CCHamide-2 (CCHa2). *CCHa2* has been suggested to be a target of Mondo, a *Drosophila* homologue of ChREBP/MondoA (Mattila et al., 2015). *CCHa2* is synthesized mainly in the fat body, an organ analogous to the mammalian liver and adipose tissues, in response to glucose ingestion (Sano et al., 2015). Fat body is the prime organ of Mondo action as *Mondo* mutant phenotype can be rescued by restoring *Mondo* only in the fat body (Havula et al., 2013). To examine whether Mondo activates *CCHa2* expression in the fat body, we knocked-down *Mondo* specifically in the fat body. The knockdown reduced not only the expression of *CCHa2* under regular culture condition (**Fig. 1A**) but also its induction upon glucose ingestion (**Fig. 1B**). This tissue-autonomous regulation by Mondo makes *CCHa2* an excellent marker for analyzing how sugars activate Mondo.

**Fig. 1.**
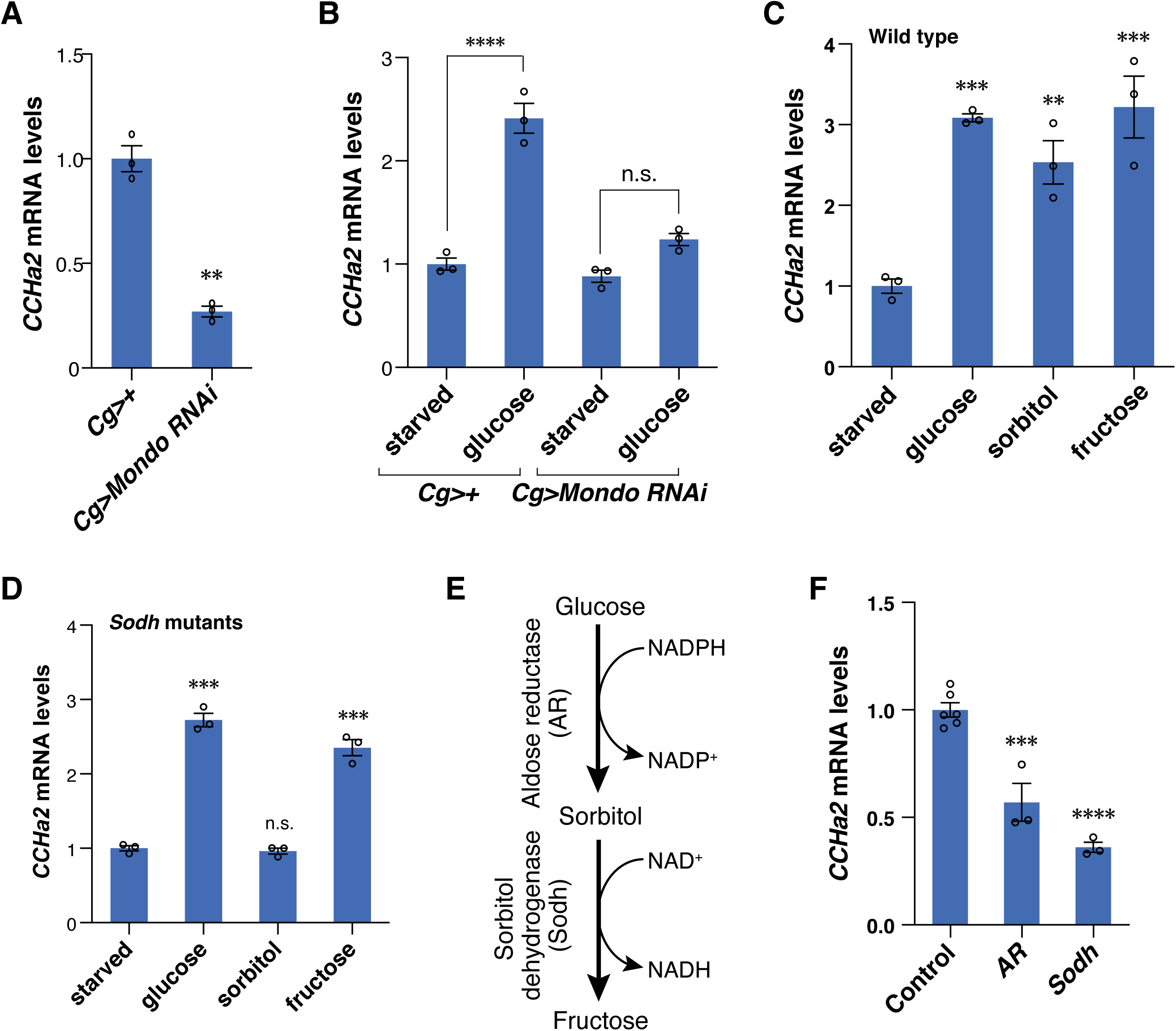
The polyol pathway is required for Mondo-mediated *CCHa2* expression. **A**, *CCHa2* mRNA levels in third-instar larvae [72 hours after egg laying (AEL)] in which *Mondo* was knocked-down in the fat body using the *Cg-GAL4* driver. **B**, *Mondo-*knockdown larvae were starved for 18 hours then re-fed with 10% glucose for 6 hours. *CCHa2* mRNA levels were quantified after re-feeding. **C**, Effects of different sugars on *CCHa2* expression. Starved wild-type larvae were re-fed for 6 hours with a 10% solution of the indicated sugars. **D**, Effects of different sugars on *CCHa2* expression in *Sodh* mutant larvae. **E**, The polyol pathway. **F**, *CCHa2* mRNA levels in the third-instar larvae of polyol pathway mutants raised on a normal diet containing 10% glucose. 10 larvae per batch, n=3 batches for all experiments. Histograms show mean ± SE. n.s., *P* > 0.05; \*\**P* < 0.01; \*\*\**P* < 0.001; \*\*\*\**P* < 0.0001.

To identify metabolic pathways required for *CCHa2* expression, we examined the effects of sugars on *CCHa2* expression. We starved *Drosophila* larvae for 18 hours then refed them with several sugars. In addition to glucose and fructose (Sano, 2015; Sano et al., 2015), sorbitol was found to be capable of inducing *CCHa2* expression (**Fig. 1C**). Because sorbitol is generated and metabolized exclusively by the polyol pathway [The Kyoto Encyclopedia of Genes and Genomics (KEGG) pathway database] (Kanehisa and Goto, 2000; Kanehisa et al., 2019), the induction of *CCHa2* expression likely involves metabolic reactions through the polyol pathway. In this pathway glucose is converted to sorbitol by aldose reductase (AR, EC: 1.1.1.21), and then to fructose by sorbitol dehydrogenase (Sodh, EC: 1.1.1.14) (**Fig. 1E**) (Kanehisa and Goto, 2000; Kanehisa et al., 2019). While AR and Sodh are also predicted to transform xylose to xylulose via xylitol (The KEGG pathway database; **Fig. S4B**) (Kanehisa and Goto, 2000; Kanehisa et al., 2019), xylitol did not induce *CCHa2* expression significantly in wild-type larvae (**Fig. S4C**). We thus reasoned that the role of the polyol pathway can be revealed by analyzing the requirement of AR and Sodh.

To create genetic tools to block the polyol pathway, we doubly mutated the putative *AR* genes, *CG6084* and *CG10638*, hereafter named *AR* mutants (**Fig. S1, S2**). We also mutated sorbitol dehydrogenase (*Sodh*) genes, *Sodh-1* and *Sodh-2*, creating what we hereafter refer to as *Sodh* mutants (**Fig. S3**). Both *AR* and *Sodh* mutants were viable and fertile but showed metabolic phenotypes as predicted (**Fig. S4**). In the mutants raised on normal fly food containing 10% glucose, *CCHa2* mRNA levels were significantly reduced (**Fig. 1F**), even though genes involved in glycolysis and PPP were intact in these animals. Thus, the polyol pathway appears to possess an independent function distinct from major glycolytic pathways. Interestingly, under starved conditions in which storage sugars were depleted (Matsuda et al., 2015), both glucose and fructose were effective in restoring *CCHa2* mRNA levels in *Sodh* mutant larvae (**Fig. 1D**). These results suggest that the polyol pathway and glycolysis/PPP have differential requirements in different nutritional conditions; the polyol pathway is dispensable for *CCHa2* expression when glucose is provided after starvation but is required for regulating its expression under normal nutritional conditions.

### The polyol pathway is required for proper larval growth and physiology

We also examined whether the polyol pathway has any physiological function in *Drosophila* larvae. The polyol pathway mutants exhibited marked loss of body weight, abnormal triacylglycerol accumulation and hemolymph glucose levels (**Fig. 2A-C**). The phenotypic difference between *AR* and *Sodh* mutants is likely caused by AR’s involvement in metabolic pathways other than the polyol pathway (Kanehisa and Goto, 2000; Kanehisa et al., 2019).

**Fig. 2.**
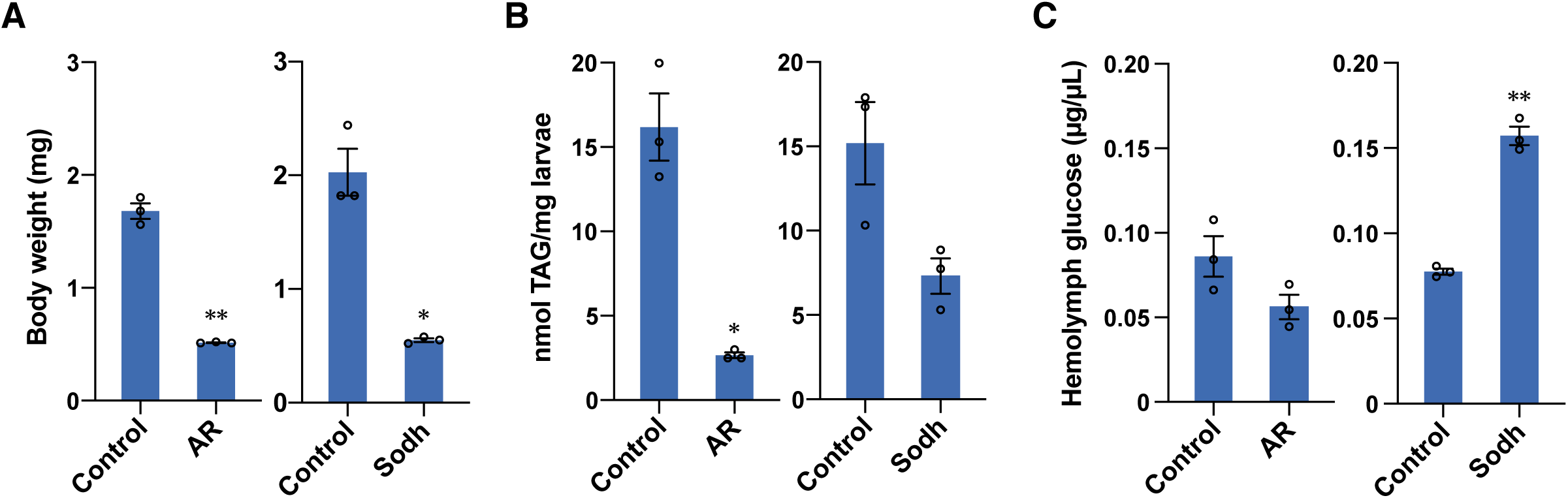
The polyol pathway is required for proper larval growth and physiology. **A**-**C**, Physiological phenotypes of *AR* and *Sodh* mutants. Body weight (**A**), triacylglycerol levels in whole body (**B**), and hemolymph glucose levels (**C**) in the third-instar larvae of control, *AR* mutant, and *Sodh* mutant were measured. The phenotypic difference between *AR* and *Sodh* mutants is likely caused by AR’s involvement in metabolic pathways other than the polyol pathway (Kanehisa and Goto, 2000; Kanehisa et al., 2019). 5-10 larvae per batch, n=3 batches. Histograms show mean ± SE. \**P* < 0.05, \*\**P* < 0.01.

### The polyol pathway is required for global transcriptional alteration and metabolic remodeling by sugar feeding

The phenotypes of *AR* and *Sodh* mutants suggest that the polyol pathway regulates not only *CCHa2* but also a wide range of Mondo/Mlx-target genes. We thus tested whether the polyol pathway couples glucose ingestion to global transcriptional alteration through Mondo. Starved larvae were re-fed with either glucose or sorbitol, and expression levels of sugar-responsive Mondo/Mlx-target genes (Mattila et al., 2015) (**Data S1**) were quantified by RNA-seq analysis. Given that sorbitol is metabolized only through the polyol pathway, polyol pathway metabolites would be selectively increased in sorbitol-fed larvae, whereas metabolites of polyol, glycolytic and PPP pathways would be increased in glucose-fed larvae. We detected a strong correlation between the changes induced by glucose and sorbitol (**Fig. 3A**). Transcriptome changes upon sorbitol feeding were lost in the *Sodh* mutants (**Fig. 3B**), confirming that sorbitol-induced gene regulation observed in wild type is dependent on the polyol pathway. Fructose, the end product of the polyol pathway restored gene regulation in the *Sodh* mutants (compare **Fig. 3C, D**), although the possibility that the influx of fructose into glycolysis participates in the rescue cannot formally be ruled out. These results show that the polyol pathway alone can regulate the great majority of Mondo/Mlx-target genes, thus playing an essential role in Mondo-mediated transcriptional regulation in response to glucose ingestion.

**Fig. 3.**
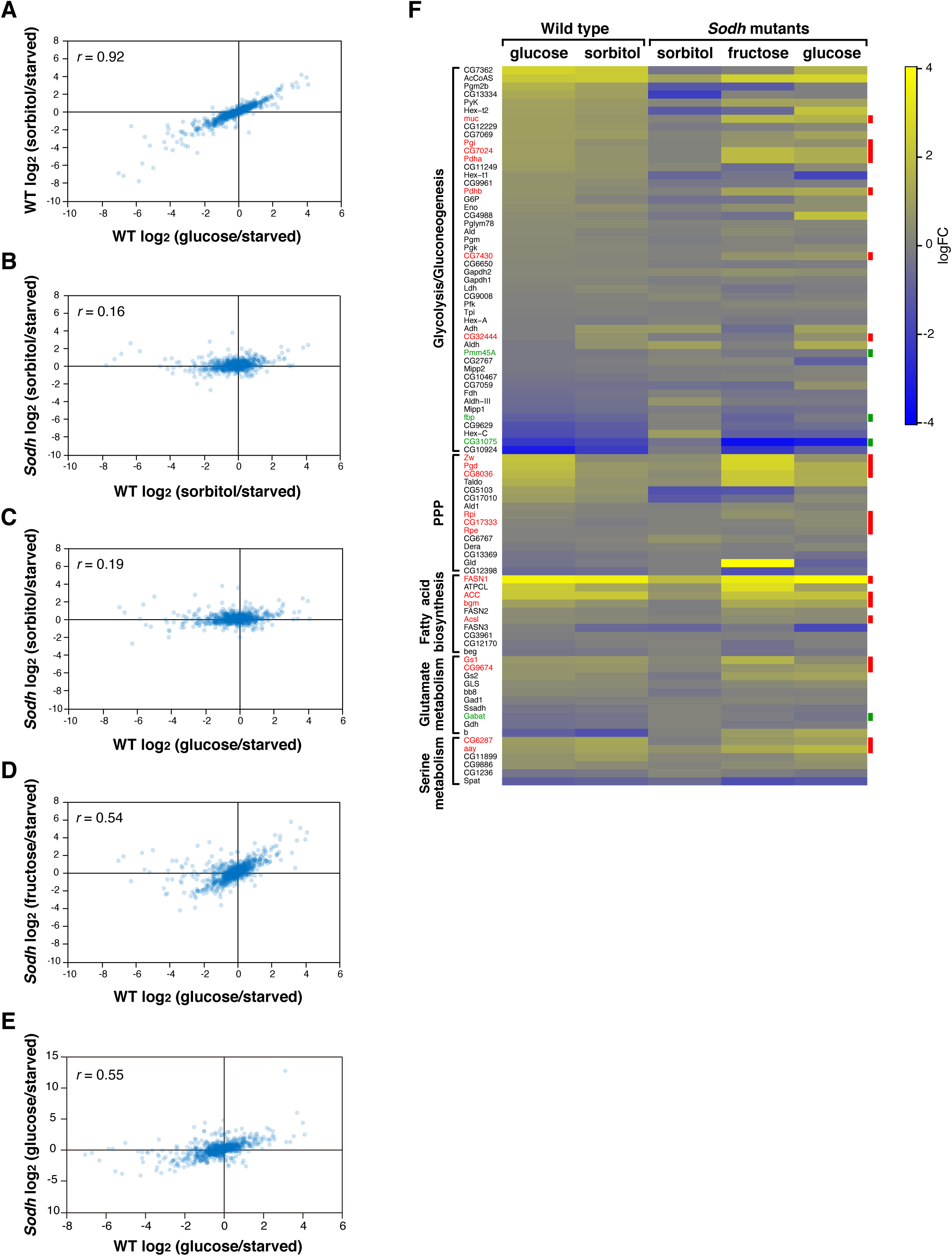
The polyol pathway is crucial for sugar-induced global transcriptional alteration. Wild-type or *Sodh* mutant third-instar larvae were starved for 18 hours, followed by re-feeding for 6 hours with a 10% solution of indicated sugars. **A-E**, A comparison of expression changes of the Mondo/Mlx-target genes between glucose-fed and sorbitol-fed wild-type larvae (**A**), sorbitol-fed wild-type and *Sodh* mutant larvae (**B**), glucose-fed wild-type and sorbitol-fed *Sodh* mutant larvae (**C**), glucose-fed wild-type and fructose-fed *Sodh* mutant larvae (**D**), glucose-fed wild-type and *Sodh* mutant larvae (**E**). **F**, The expression changes of metabolic genes. Genotype of larvae and fed sugars are indicated above. Known Mondo target genes are indicated in red (activated genes) or green (suppressed genes). 30 larvae per batch, n = 3 batches. Correlation coefficients (*r)* are indicated in the plots (**A-E**).

We then examined whether the polyol pathway triggers Mondo-mediated metabolic remodeling. We focused on genes encoding enzymes involved in glycolysis/gluconeogenesis, PPP, fatty acid biosynthesis, and glutamate and serine metabolism, many of which are under the control of Mondo (Mattila et al., 2015). We observed that feeding of glucose and sorbitol caused similar changes in the expression patterns of these metabolic genes in wild-type larvae (**Fig. 2F**). Changes in metabolic gene expression were reduced in sorbitol-fed *Sodh* mutant larvae, but were restored in *Sodh* mutant larvae when fructose was fed. In particular, the levels of known Mondo/Mlx-target genes were remarkably restored (indicated in red and green in **Fig. 2F**). These results show that the polyol pathway is crucial for the expression of various metabolic enzymes, leading to a metabolic remodeling in response to sugar ingestion. Additionally, when starved *Sodh* mutant animals were fed with glucose, a considerable number of Mondo/Mlx-target genes were regulated properly (**Fig. 3E, F**). These results, together with the data shown in Figure 1D, suggest that glucose-metabolizing pathways other than the polyol pathway can activate Mondo under starved conditions, in which glycogen is completely consumed in the fat body (Matsuda et al., 2015). In such situation, the activity of glycolysis and PPP could also reflect glucose uptake and function as a glucose sensor leading to Mondo activation (see discussion).

### The polyol pathway regulates nuclear localization of Mondo

The above results suggest that the polyol pathway is involved in a critical step in the regulation of Mondo under normal physiological conditions. Therefore, we examined the effects of polyol pathway mutations on nuclear localization of Mondo. We tagged endogenous Mondo with the Venus fluorescent protein (**Fig. S5**), and observed intracellular localization of the Mondo::Venus fusion protein in *ex vivo* culture of fat bodies dissected from normally-fed third-instar larvae. It has been reported that mammalian ChREBP/MondoA displays nuclear localization when glucose concentrations are increased five to ten-fold (Arden et al., 2012; Dentin et al., 2012; Kabashima et al., 2003; Kawaguchi et al., 2001; Li et al., 2006; Noordeen et al., 2012; Petrie et al., 2013; Sakiyama et al., 2008). Therefore, we compared the nuclear localization of the Mondo::Venus protein in fat bodies cultured in Schneider’s *Drosophila* medium (hereafter referred to as the basic medium) that contains 11 mM glucose and those cultured in the same medium supplemented with 55 mM sugars. In the basic medium, 5.2% of Mondo::Venus signals were localized in the nuclei of wild-type fat body cells (**Fig. 4A, B**). When glucose, sorbitol, or fructose were added to the basic medium, the percentage of nuclear Mondo::Venus signals was increased to 14.6%, 12.7%, and 16.7%, respectively (**Fig. 4A, B**). In contrast, in the *AR* mutant or *Sodh* mutant fat body cells, glucose administration did not increase nuclear Mondo::Venus signals, suggesting that the polyol pathway required for the activation of Mondo (**Fig. 4A, C, D**). Indeed, metabolites generated in the polyol pathway bypassed the requirements for *AR* or *Sodh* in the nuclear localization of Mondo under fed conditions (**Fig. 4A, C, D**). Taken together, we propose that the polyol pathway acts as a system for sensing glucose uptake that allows metabolic remodeling.

**Fig. 4.**
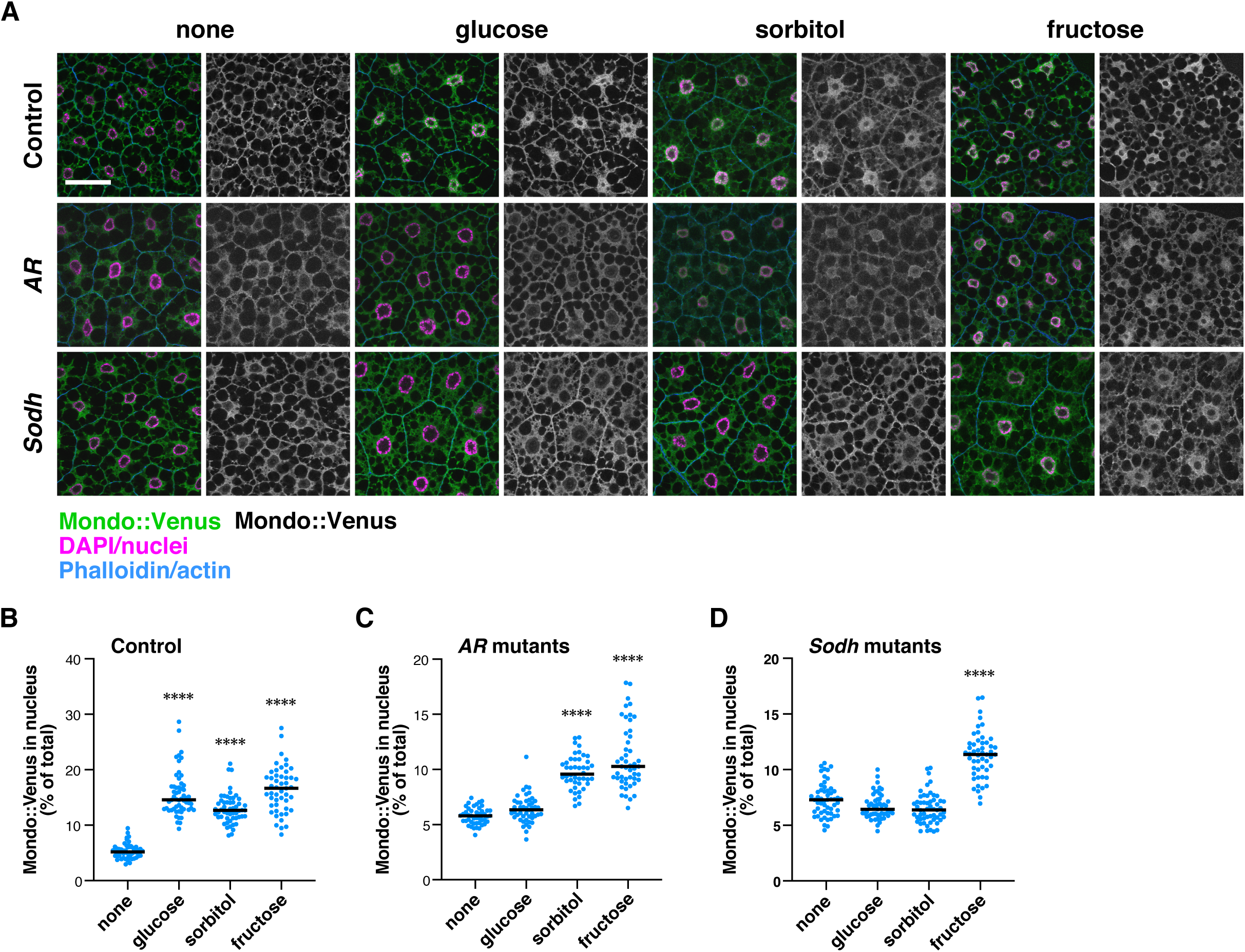
The polyol pathway regulates nuclear localization of Mondo. **A**, Fat bodies dissected from the Mondo::Venus knockin line were cultured in Schneider’s *Drosophila* medium supplemented with 55 mM glucose, sorbitol, or fructose for 15 minutes. After culture, the fat bodies were fixed and stained with the following markers: anti-GFP antibody for Mondo::Venus (green), DAPI (magenta), and Rhodamine-conjugated phalloidin (blue). **B-D**, The percentage of nuclear Mondo::Venus signal out of the total Mondo::Venus signal in a cell was quantified in the images. Images of 44 to 56 fat body cells per experiment were quantified. Black horizontal lines show the median Scale bar represents 50 µm. \*\*\*\**P* < 0.0001.

### The polyol pathway also functions as a glucose-sensing system in mouse liver

To clarify whether the function of the polyol pathway in the sensing of glucose uptake is evolutionarily conserved, we investigated the nuclear localization of ChREBP in response to sugar ingestion in mouse liver. To remove ChREBP from the nuclei of the hepatocytes, we starved mice overnight. We then orally administered a sugar solution to starved mice and examined the intracellular localization of ChREBP in the hepatocytes. We focused on pericentral hepatocytes as Sorbitol dehydrogenase (Sord), the only enzyme catalyzing the second step of the polyol pathway in mice, is preferentially expressed (Halpern et al., 2017; LeCluyse et al., 2012). Glucose or fructose ingestion promoted nuclear localization of ChREBP in wild-type mice (**Fig. 5A-C, G-I, M**). To examine whether glucose-responsive nuclear translocation of ChREBP is mediated by the polyol pathway, we generated *Sord* knockout mice using the CRISPR-Cas9 system (**Fig. S6B**). In *Sord* knockout mice, glucose administration did not promote nuclear translocation of ChREBP (**Fig. 5D, E, J, K, M**), whereas fructose ingestion did (**Fig. 5F, L, M**). The *Sord* knockout mice showed impaired glucose tolerance, namely a delay in the recovery of blood glucose levels after oral glucose administration (**Fig. 5N, O**). These results indicate that the polyol pathway has an important function in sensing glucose uptake in mouse liver, and its deficiency leads to impaired glucose tolerance. Thus, the polyol pathway is a common system for sensing glucose uptake in flies and mouse.

**Fig. 5.**
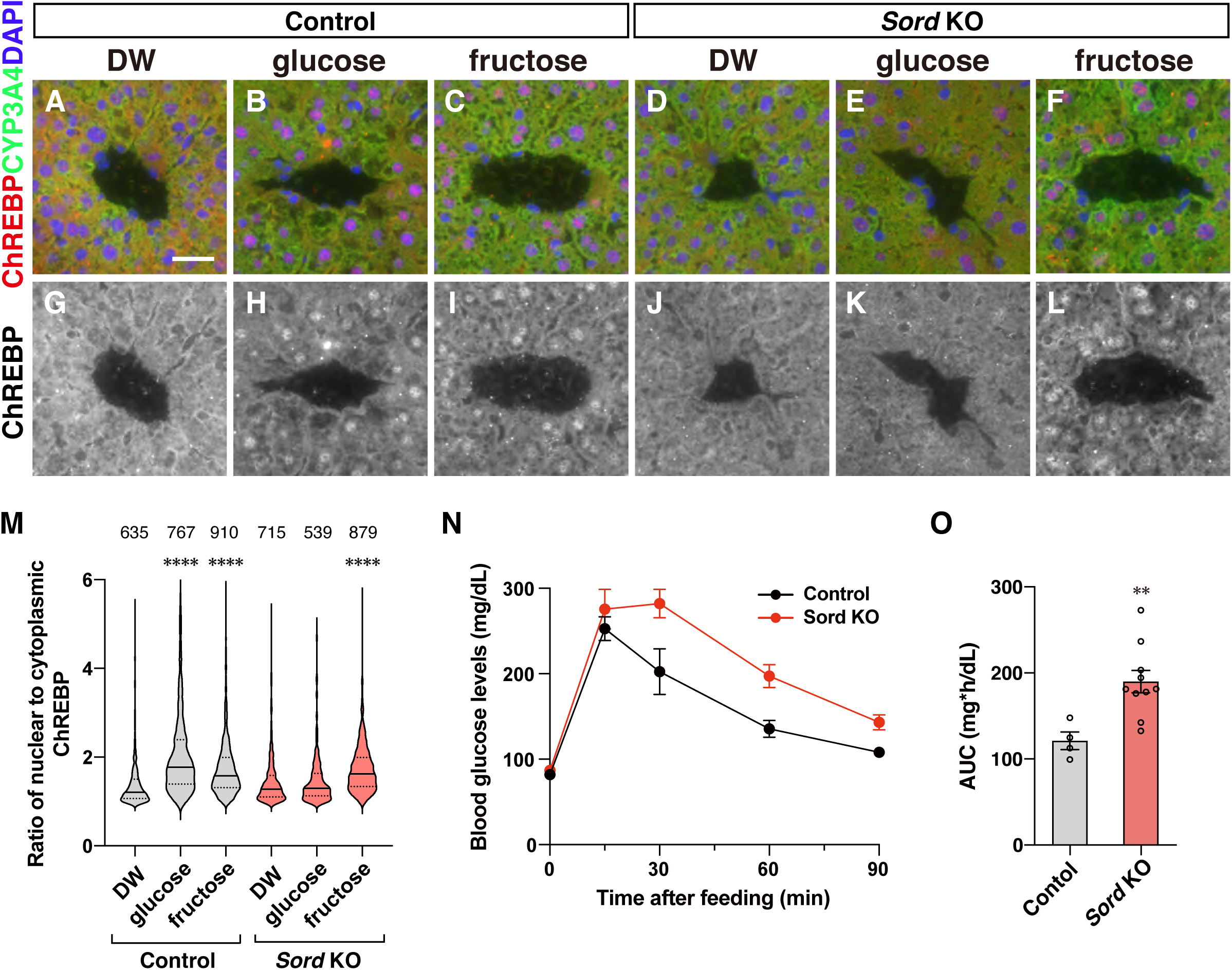
The polyol pathway regulates nuclear localization of ChREBP in hepatocytes and glucose tolerance in mice. **A-L**, Water (DW), glucose, or fructose were administered orally to starved control and *Sord* knockout mice, and the localization of ChREBP in the hepatocytes was examined 15 minutes after administration. Liver slices were stained with the following markers: anti-ChREBP antibody (red), anti-CYP3A4 for pericentral hepatocytes (green), and DAPI for nuclei (blue). Scale bar in (**A**) represents 30 µm. **M**, Ratio of average intensity of ChREBP signals detected in the nucleus and cytoplasm in the hepatocytes. Solid and dotted lines in the graph show the median and quartiles, respectively. n=3 animals. The number of cells scored are indicated on the top. **N**, Blood glucose levels were measured over a 90-minute period after glucose administration. **O**, Area under the curve (AUC) was calculated relative to the fast blood glucose concentrations. n=4 for control, n=10 for *Sord* KO. \*\**P* < 0.01; \*\*\*\**P* < 0.0001.

## Discussion

In this paper, we have shown that the polyol pathway functions as a conserved system for sensing glucose uptake. Genome research has revealed that polyol pathway enzymes are conserved from yeasts to humans, suggesting that this pathway is important across species. However, based on the low affinity of AR for glucose, it has long been believed that the polyol pathway is almost silent, and is only activated under hyperglycemic conditions, leading to diabetic complications (Gabbay, 1973). Our study revealed the evolutionarily conserved function of the polyol pathway in organismal physiology.

### The significance of the polyol pathway in sugar sensing

To enable proper organismal adaptation to ingested sugars, the activity of the metabolic pathway(s) required for sugar sensing is expected to correlate with the levels of glucose in the body fluid. The polyol pathway appears to fulfill these conditions. First, the polyol pathway is the most upstream glucose-metabolizing pathway. Glucose flows into the polyol pathway before being metabolized to glucose-6-phosphate, which is consumed through glycolysis and PPP (**Fig. 6**). Second, the polyol pathway would be less affected by storage sugars. Glycogen, the major carbohydrate storage form in animal cells, is converted reversibly into glucose-6-phosphate according to nutrient status of the cell (**Fig. 6**). The adjustments to different nutritional states maintain constant glucose-6-phosphate levels, thereby aiding the stability of glycolysis and PPP regardless of the availability of glucose (Peeters et al., 2017). Third, no feedback control on the polyol pathway has been reported. This is a sharp contrast to glycolysis, in which several enzymes are subject to feedback control by downstream metabolites. Hexokinase acting at the most upstream point in glycolysis is tightly regulated by its product (Berg, 2006). Phosphofructokinase 1, a rate-limiting enzyme of glycolysis, is also controlled by several downstream metabolites such as ATP, AMP, citrate, lactate, and fructose-2,6-bisphosphate (Mor et al., 2011). These observations suggest that the polyol pathway could exhibit a linear response to glucose levels in the body fluid better than glycolysis and PPP under normal feeding conditions. Therefore, it is conceivable that the polyol pathway acts as a glucose-sensing system under conditions in which homeostasis of major glucose metabolic pathways is maintained by storage sugars and feedback control. Our results also suggest that multiple glucose-sensing pathways exist and function in different nutritional conditions (**Fig. 1D, 3E, F**), which might be related to the identification of glycolytic and PPP-derived metabolites as ChREBP-activating sugars in mammalian cell culture systems (**Fig. 6**). Having various glucose sensing systems would be beneficial for cells and organisms for their adaptation to different types of changes in nutritional conditions.

**Fig. 6.**
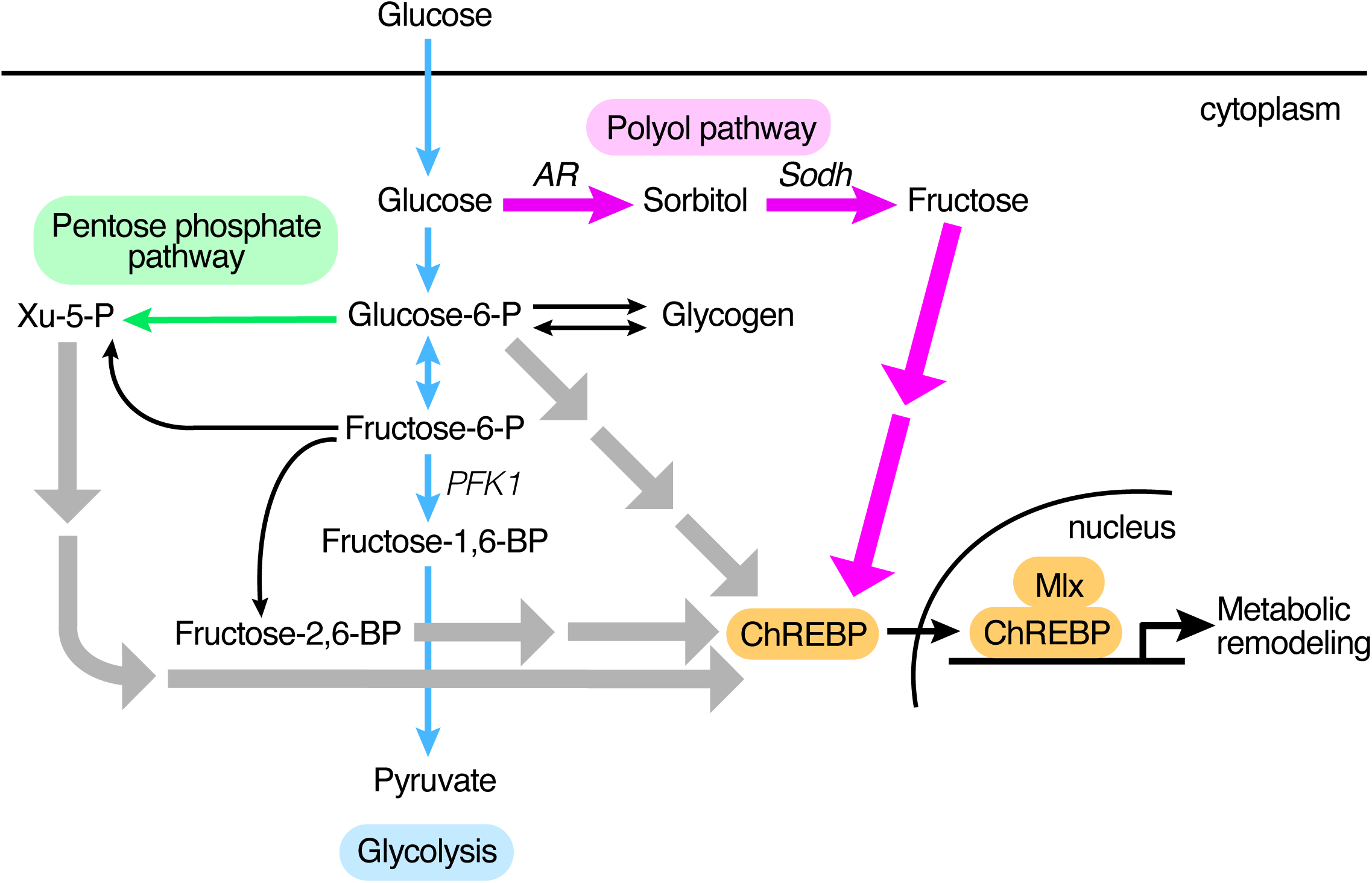
Metabolic pathways leading to Mondo activation. Glucose-6-phosphate, xylulose-5-phosphate, and fructose 2,6-biphosphate were identified previously as ChREBP-activating metabolites in mammalian cell culture systems (gray arrows) (Richards et al., 2017). These metabolites are generated during glycolysis (blue arrows) or PPP (green arrow). Our study revealed that the polyol pathway (magenta arrows) has a significant contribution to activating Mondo/ChREBP in flies and mice (magenta arrows). HK: hexokinase, PFK1: phosphofructokinase 1, PFK2: phosphofructokinase 2.

Our results suggest that fructose and fructose derivatives are good candidates for metabolite(s) that activates Mondo/ChREBP in sensing glucose uptake via the polyol pathway. Fructose appears to have cell-autonomously and cell-nonautonomous functions. In fly larvae, *AR* is expressed ubiquitously, and *Sodh* is preferentially expressed in the fat body and gut (Graveley, 2011). This is consistent with our observation that the polyol pathway functions as a glucose-sensing system in the fat body. Additionally, fructose or fructose derivatives is thought to be secreted into the hemolymph from which they signal to cells in other organs. It has been shown that the concentration of circulating fructose is acutely elevated upon glucose ingestion, probably due to the low basal concentration of fructose in the hemolymph (Miyamoto et al., 2012). Therefore, conversion of a portion of ingested glucose to fructose could be advantageous to allow glucose detection, especially in hyperglycemic animals such as insects.

Hyperglycemia is also observed in the mammalian liver to which dietary glucose is carried directly from the small intestine through the portal vein. We have shown that the polyol pathway is required for sensing glucose uptake in the mouse liver. The hepatic lobules are compartmentalized into regions with different metabolic functions along the porto-central axis: glycolysis and lipogenesis occur in the hepatocytes close to the central vein (Kietzmann, 2017). *Sord* mRNA is expressed with a peak in the pericentral hepatocytes (Halpern et al., 2017), suggesting that the polyol pathway functions in the same region where glycolysis and lipogenesis occur and contributes to matching the activities of glycolysis and lipogenesis with glucose supply. On the other hand, whether fructose is released into the circulation and signals to other cells in the liver and other organs awaits further analysis.

### Insights into fructose-induced pathogenic mechanisms

The model that fructose or fructose derivatives activate Mondo/ChREBP can explains the beneficial as well as harmful effects of fructose. We have shown that the polyol pathway, i.e., the presence of fructose, decreases the glycemic responses to oral glucose intake in mice (**Fig. 5N, O**). Consistently, it has been shown that a small amount of fructose improves glucose tolerance in healthy and diabetic adults (Crapo et al., 1980; Moore et al., 2000; Moore et al., 2001). Our results suggest that metabolic remodeling governed by the polyol pathway accounts for these phenomena. On the other hand, it is well known that excessive fructose intake, as represented by high-fructose corn syrup, has adverse effects on human health (Bray et al., 2004; Lim et al., 2010; Marriott et al., 2009). High-fructose corn syrup has been used in artificially sweetened foods since the 1970s, and fructose consumption has increased drastically over the past decades. Epidemiological studies have shown that the increase in fructose consumption correlates with that in metabolic diseases including obesity, fatty liver, and nonalcoholic fatty liver disease (Bray et al., 2004; Lim et al., 2010; Marriott et al., 2009). Experimental studies have revealed that fructose administration to the cell elevates lipid accumulation better than glucose does (Stanhope et al., 2009; Theytaz et al., 2014). It has been proposed that fructose is harmful because it is converted to fructose-1-phosphate by fructokinase, which accelerates glycolysis by evading the rate-limiting steps of glycolysis and promotes lipogenesis by generating dihydroxyacetone phosphate (Heinz et al., 1968). Although this appeared plausible when fructose was believed to be metabolized in the liver, it became less credible as fructose was shown to be cleared in the small intestine by ketohexokinase (Jang et al., 2018). Although overconsumption of fructose causes its leakage to the liver (Jang et al., 2018), such small amount of fructose would not fully account for its metabolic toxicity. Our results provide an alternative explanation for the toxicity of fructose. Glucose is a poor substrate for the polyol pathway as its Km value for AR is 70 to 150 mM (Gabbay, 1973). Therefore, only very small amounts of fructose would be converted from glucose through the polyol pathway under normal feeding conditions. A direct inflow of fructose to the liver could mislead the cells into responding as if there has been very high level of glucose consumption, causing them to over-activate metabolic responses through ChREBP. Consistent with this, high-fructose ingestion in mice and rats is associated with increased ChREBP activity in the liver (Kim et al., 2016; Koo et al., 2009). Therefore, it is likely that high-fructose corn syrup is particularly deleterious to human health because it triggers drastic metabolic remodeling through ChREBP in the liver.

Fructose is also implicated in cancer development. Feeding mice with high-fructose corn syrup enhances tumor growth independently of obesity and metabolic syndrome (Goncalves et al., 2019). Interestingly, expression levels of the polyol pathway enzyme *AR* correlate with the epithelial-to-mesenchymal transition (EMT) status in cancer cell lines as well as in cancers in patients. Moreover, knockdown of *AR* or sorbitol dehydrogenase (*Sord)* genes can block EMT *in vitro* (Schwab et al., 2018). These observations suggest that the polyol pathway links sugar metabolism to cancer metastasis. Our work lays the foundation for further important studies uncovering the molecular mechanisms linking abnormal sugar metabolism and disease development.

## Acknowledgements

We thank Shu Kondo, the Bloomington Drosophila Stock Center, the Vienna Drosophila Resource Center, FlyORF in University of Zurich for supplying plasmid and fly stocks. We also acknowledge Masayo Yamane, Miyuki Nishigata, Takumi Ichikawa, Shingo Usuki, Masatake Araki, Yoko Kimachi, Narumi Koga, and Yuki Takada for technical assistance, and Daria Siekhaus, Prashanth Rangan, Takashi Koyama, and Yasushi Hiromi for critical reading of the manuscript.

## Author Contributions

Conceptualization: HS

Investigation: HS, MY, HN, TN, KA, KT, KI, HA

Writing-original draft: HS

Writing-review & editing: HS, AN, HA

Funding acquisition: HS, AN, MK

Resources: HS, AN

Supervision: HS, AN

## Declaration of Interests

The authors declare no competing interests.

## Data availability

All datasets generated during this study are available on NCBI GEO (accession #).

## Materials and Methods

### Fry strains and dietary conditions

The following fly stocks were used: *Oregon-R (OR), white (w), y w, Cg-GAL4* (BDSC, RRID: BDSC_7011), *UAS-Mondo RNAi* (VDRC, v109821). *CG6084*^*10-1*^, *CG10638*^*10-1*^, *sodh1*^*14-1*^, and *sodh1*^*9-3*^ were generated using the CRISPR/Cas9 system (see below). Flies were raised at 25°C on regular fly food containing (per liter) 40 g yeast extract, 50 g cornmeal, 30 g rice bran, 100 g glucose, and 6 g agar.

### Mutagenesis and Venus knockin

The polyol mutants were generated using the CRISPR/Cas9 system as described in Gokcezade *et al*. (2014) (Gokcezade et al., 2014). The following sgRNA targets were used for the mutagenesis of the genes encoding AR and Sodh. Breakpoints of the mutants were determined as described previously (Kina et al., 2019; Sano et al., 2015) (**Fig. S1-S3**). *CG6084* and *CG10638* were doubly mutated in *AR* mutants. *Sodh-1* and *Sodh-2* were doubly mutated in *Sodh* mutants.

*CG6084*: 5′-CCCCAAGGGTCAGGTCACCG

*CG10638*: 5′-GGCTACGAGATGCCAATTCT

*Sodh-1*: 5′-GATGTACACTACCTTGCACA

*Sodh-2*: 5′-GTGGGCAAGGTAGTGCACGT

The knockin of Venus at the C-terminus of the Mondo coding region was performed using the CRISPR/Cas9 system. The knockin vector was constructed by combining PCR-amplified left arm and right arm fragments for homologous recombination, and the Esp3I fragment of the pPVxRF3 vector (a gift from S. Kondo) containing Venus and 3xP3-dsRed-Express2 using NEBuilder HiFi DNA Assembly Master Mix (NEB). The combined fragment was cloned into pBluescript. The oligonucleotides used are as follows:

L-arm forward: 5′-GCTTGATATCGAATTCTGAACGACTGGAAATTTTGG

L-arm reverse: 5′-AGTTGGGGGCGTAGGGGGTGCATGCAGATTTGG

R-arm forward: 5′-TAGTATAGGAACTTCCGTTGATGCTGATGTCCTTG

R-arm reverse: 5′-CGGGCTGCAGGAATTCGAAAATGAGAGAAGATGGCGTA

The knockin vector was injected into *y w* embryo together with the sgRNA plasmid. The following sgRNA target was used.

5′-GGCCAGCATCCAAATCTGCA

### Quantitative RT-PCR

Quantitative RT-PCR was performed as described previously(Sano et al., 2015). The following primers were used:

*CCHa2* forward: 5′-GCCTACGGTCATGTGTGCTAC

*CCHa2* reverse: 5′-ATCATGGGCAGTAGGCCATT

*rp49* forward: 5′-AGTATCTGATGCCCAACATCG

*rp49* reverse: 5′-CAATCTCCTTGCGCTTCTTG

### RNA-sequencing

Third-instar larvae (72 hours AEL) were starved for 18 hours on water agar plates. The larvae were re-fed on agar plates containing 10% of indicated sugar for 6 hours. Sugar plates were supplemented with 1% Brilliant Blue to visualize larval sugar ingestion. Total RNA from whole larvae was extracted using the PureLink RNA Mini Kit (Life Technologies). The library was constructed using the TruSeq Stranded mRNA LT Sample Prerp Kit (Illmina). RNA-seq was performed with NextSeq 500 (Illumina), targeting at least 14 million, single-end reads of 75 bp in size. The quality of the reads was assessed using FastQC (version 0.11.5). The reads were mapped to the FlyBase reference genome (Dmel Release 6.19) using Tophat2(Kim et al., 2013). Transcript abundance and splice variant identification were determined using Cufflinks(Trapnell et al., 2010), and differential expression analysis was performed using CuffDiff(Trapnell et al., 2010). We confirmed that gene expression patterns correlate well in starved wild-type and *Sodh* mutant larvae, indicating that the observed differences in sugar-dependent gene expression between wild type and mutants are not due to variations in their genetic background (**Fig. S7**).

### Measurement of triacylglycerol and glucose

For measurement of triacylglycerol (TAG) concentration in the whole body, third-instar larvae (96 hours AEL) were homogenized in water with NP-40. The homogenate was heated at 90°C for 5 minutes and then mixed by vortex, which was repeated twice. The homogenate was centrifuged at 16,000 *g* for 2 minutes, and the supernatant was used for TAG quantification using the Triglyceride Quantification Colorimetric/Fluorometric Kit (BioVision). For measurement of hemolymph glucose levels, third-instar larvae (96 hours AEL) were rinsed with water, and dried on filter paper. The cuticle was torn by forceps to release the hemolymph on a Parafilm membrane. 1 µl of hemolymph was diluted with 9 µL of Tris buffered saline (ph. 6.6) and immediately heated at 70°C for 5 minutes. The hemolymph solution was centrifuged at 16,000 *g* for 1 min, and the supernatant was used for glucose quantification using the Glucose Colorimetric/Fluorometric Assay Kit (BioVision).

### Metabolic assays using gas chromatography – mass spectrometry

Third-instar larvae (96 hours AEL) were rinsed with water, and dried on filter paper. The cuticle was torn by forceps to release the hemolymph on a Parafilm membrane. 1 µl of hemolymph was collected and immediately quenched by mixing with 300 µl of cold methanol. The samples were further mixed with 200 µl of methanol, 200 µl of H_2_O, and 200 µl of CHCl_3_, and vortexed for 20 min at room temperature. The samples were centrifuged at 20,000 *g* for 15 min at 4°C. The supernatant was mixed with 350 µl of H_2_O and vortexed for 10 min at room temperature. The samples were centrifuged at 20,000 *g* for 15 min at 4°C. The aqueous phase was collected and dried in a vacuum concentrator. Methoxyamine pyridine solution [20 mg/ml methoxyamine hydrochloride (Wako) in pyridine] was added to the dried residue to re-dissolve and oximated for 90 min at 30°C. Then, MSTFA + 1%TMCS (Thermo) was added and incubated for 60 min at 37°C for trimethylsilylation. The derivatized metabolites were analyzed by an Agilent 7890B GC coupled to a 5977A Mass Selective Detector (Agilent Technologies) under the following conditions: carrier gas, helium; flow rate, 0.8 ml/min; column, DB-5MS + DG (30 m × 0.25 mm, 0.25 µm film thickness; Agilent Technologies); injection mode, 1:10 split; inlet temperature, 250°C; ion source temperature, 230°C; quadrupole temperature, 150°C. The column temperature was held at 60°C for 1 min, and then increased to 325°C at a rate of 10°C/min. The detector was operated in the electron impact ionization mode. The Agilent-Fiehn GC/MS Metabolomics RTL Library was used for metabolite identification(Kind et al., 2009). Metabolites were detected in SIM mode and the peak area of interests were analyzed by the QuantAnalysis software (Agilent Technologies).

### Culture of larval fat body

In order to observe the nuclear localization of Mondo with minimal effects of larval feeding conditions and dissection, we used an *ex vivo* culture system of larval fat bodies. Fat bodies were dissected and cultured in Schneider’s *Drosophila* medium, referred to as basic medium, that contains 11 mM glucose. It has been reported that mammalian ChREBP displays nuclear localization when glucose concentrations are increased five to ten-fold (Kawaguchi et al., 2001; Li et al., 2006; Noordeen et al., 2012; Petrie et al., 2013; Sakiyama et al., 2008). Therefore, we compared the nuclear localization of Mondo::Venus in fat bodies cultured in the basic medium and in fat bodies cultured in the same medium supplemented with 55 mM glucose, sorbitol or fructose.

### Immunofluorescence and image analysis of fat bodies

Larval fat bodies were fixed with 4% paraformaldehyde in PBS for 30 minutes. Mondo::Venus was detected with rabbit anti-GFP polyclonal antibody (Thermo Fisher Scientific, 1: 1,000) and Alexa Fluor 488-conjugated anti-rabbit-IgG (Thermo Fisher Scientific, 1: 500). Nuclei and cortical actin were labelled with DAPI (Thermo Fisher Scientific, 1 µg/mL) and Rhodamine-conjugated Phalloidin (Thermo Fisher Scientific, 1: 100), respectively. After staining, fat bodies were mounted in VECTASHIELD Mounting Medium (Vector Laboratories) and imaged with TCS SP8 confocal microscope using a Plan-Apochromat 63x oil-immersion objective lens (Leica Microsystems) or Fluoview FV1000 confocal microscope using a UPlanSApo 60x water-immersion objective lens (Olympus). Images of the fat body were analyzed using the ImageJ2 software (version 2.0.0-rc-43, NIH).

### Generation of *Sord* knockout mouse

*Sord* knockout mice were generated as described previously by introducing the Cas9 protein, tracrRNA, crRNA and ssODN into C57BL/6N fertilized eggs (Takemoto et al., 2020). For generating the *Sord Δex3-9* allele, the synthetic crRNA was designed to direct GAGACAAAGGAAACACGTGA(GGG) in the intron 2 and AATCACAGTAGAACACACAA(AGG) in the exon 9. ssODN: 5’-TTCTTCATAAGTCAGCCCCACTCTCTGGCAATCACAGTAGTTTATTTATTTATGAGGGAAAGGCGAACCTTCCATTGCTCTCAGAAGTGCTA was used as a template for homologous recombination. The genome of targeted F0 mice was amplified by PCR using the Sord 13357- and Sord -30158 primers. A 1052 bp fragment was amplified from the genome of the *Sord Δex3-9* allele. The PCR amplicons were sequenced using the Sord 13815-primer. F0 mice were backcrossed with C57BL/6N to establish the *Sord Δex3-9* line.

Sord 13357-: 5’-GCAGTCTCTGGCCAGTTTTC

Sord -30158: 5’-TTGCCTGTGAGTGACTCTGG

Sord 13815-: 5’-CGGTTTCCTTTGGAATCTCA

### Oral administration of sugars in mice

Eight-week-old male mice were starved for 16 hours before the experiment. Mice were weighed and given a sugar solution (20% glucose or 30% fructose in water) of 10 µL per gram of body weight using a plastic feeding needle (1.18 × 38 mm). Blood glucose levels were measured before and after administration of the sugar solution. The animals were euthanized 15 minutes after the sugar administration. The liver was removed, embedded in the Tissue-Tek O.C.T. compound (Sakura Finetek) and frozen in liquid nitrogen-cooled isopentane.

### Oral glucose tolerance test

Eight-week-old male mice were starved for 16 hours before the experiment. Mice were weighed and given a 20% glucose sugar solution of 10 µL per gram of body weight using a plastic feeding needle (1.18 × 38 mm). Blood glucose levels were measured over a 90-minutes period after glucose administration. Area under curve (AUC) was calculated relative to the fasted blood glucose concentration.

### Immunofluorescence and image analysis of mouse hepatocytes

Immunofluorescence staining was performed on 5-µm frozen section of the liver. The frozen sections were fixed with 4% paraformaldehyde in PBS for 10 minutes at 4°C. ChREBP was detected with rabbit anti-ChREBP polyclonal antibody (Novus Biologicals, 1:100) and Cy3-conjugated anti-rabbit-IgG (Jackson ImmunoResearch, 1:500). Pericentral hepatocytes were labelled with mouse-anti-CYP3A4 antibody (Proteintech Group, 1:300) and Alexa Fluor 488-conjugated anti-mouse-IgG (Thermo Fisher Scientific, 1:500). Nuclei were labelled with DAPI (Thermo Fisher Scientific, 1 µg/mL). After staining, the sections were mounted in VECTASHIELD HardSet Antifade Mounting Medium (Vector Laboratories) and imaged with Biorevo BZ-9000 fluorescence microscope using a Plan-Apochromat 40x objective lens (Keyence). Image analysis was performed using ArrayScan XTI (Thermo Fisher Scientific) and the FlowJo 10 software (Becton Dickinson). ChREBP signals in CYP3A4 positive hepatocytes were quantified.

### Statistics

Two-tailed t-test was used to evaluate the significance of the results between two samples. For multiple comparisons, Tukey-Kramer or Dunnett test was used. A p-value of less than 0.05 was considered statistically significant.

## Supplemental Information

**Fig. S1.**
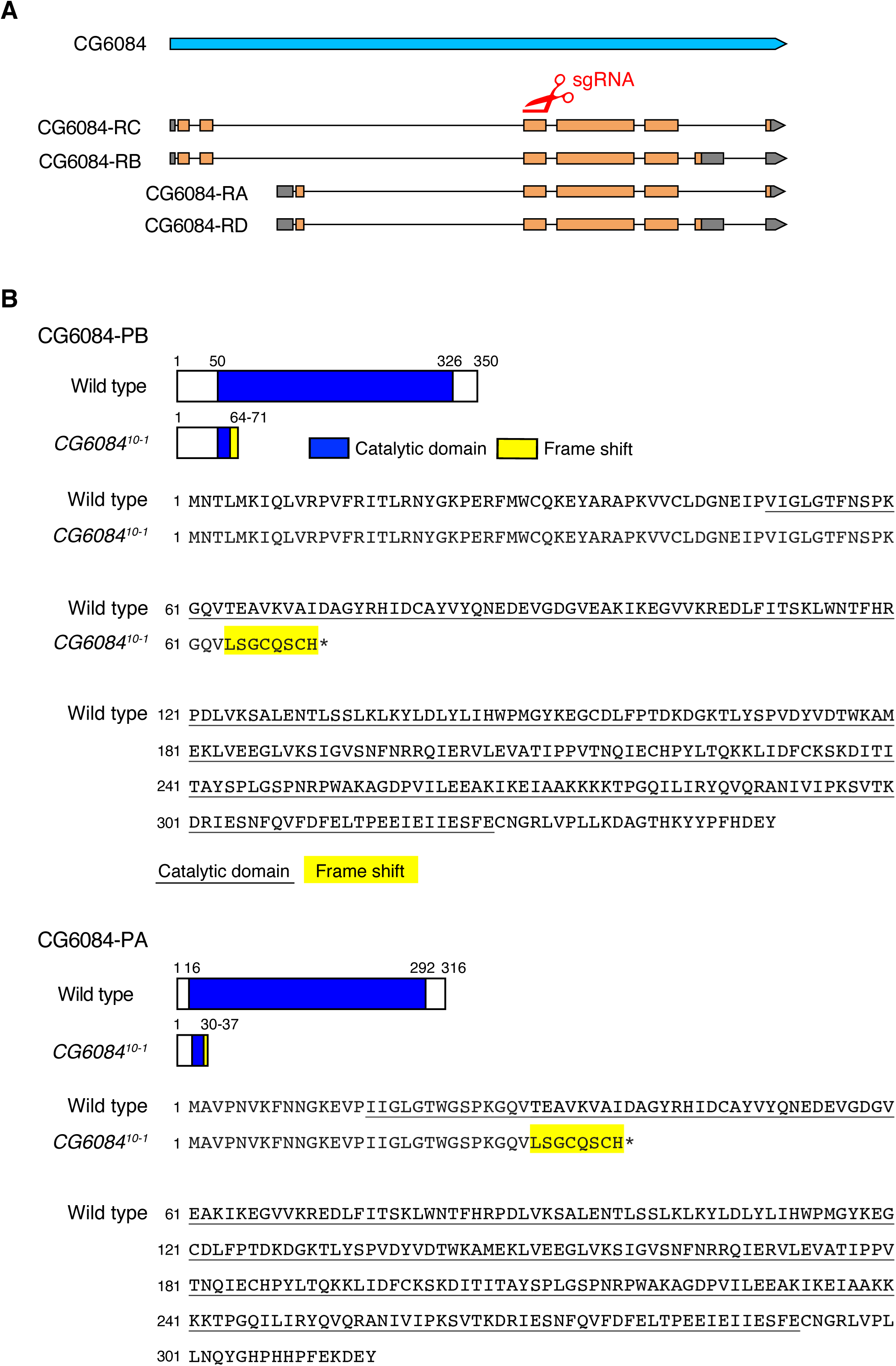
Generation of the *CG6084* mutant allele. **A**, CRISPR-mediated mutagenesis of *CG6084*. A sgRNA was designed against the sequence in the exon common to the *CG6084* isoforms. **B**, Breakpoint of the *CG6084*^*10-1*^ allele. The *CG6084*^*10-1*^ mutation caused a frame-shift (yellow) leading to a premature termination in all isoforms of the CG6084 protein. The mutant proteins lack most of the catalytic domain of the CG6084 protein (blue in schematic, underlined in sequence).

**Fig. S2.**
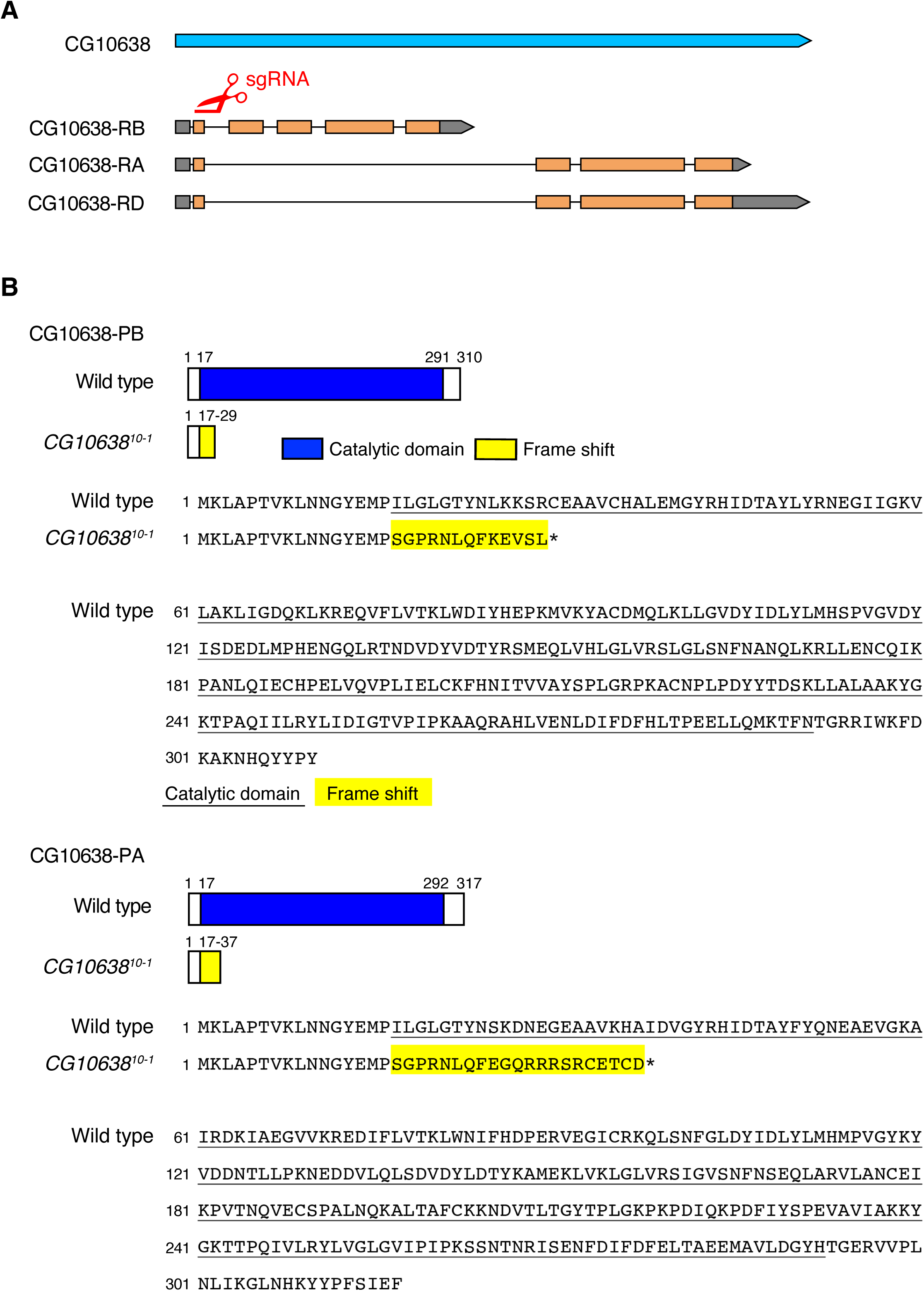
Generation of the *CG10638* mutant allele. **A**, CRISPR-mediated mutagenesis of *CG10638*. A sgRNA was designed against the sequence in the common exon of the *CG10638* isoforms. **B**, Breakpoint of the *CG10638*^*10-1*^ allele. The *CG10638*^*10-1*^ mutation caused a frame-shift (yellow) leading to premature termination of all isoforms of the CG10638 protein. The mutant proteins lack most of the catalytic domain (blue in schematic, underlined in sequence).

**Fig. S3.**
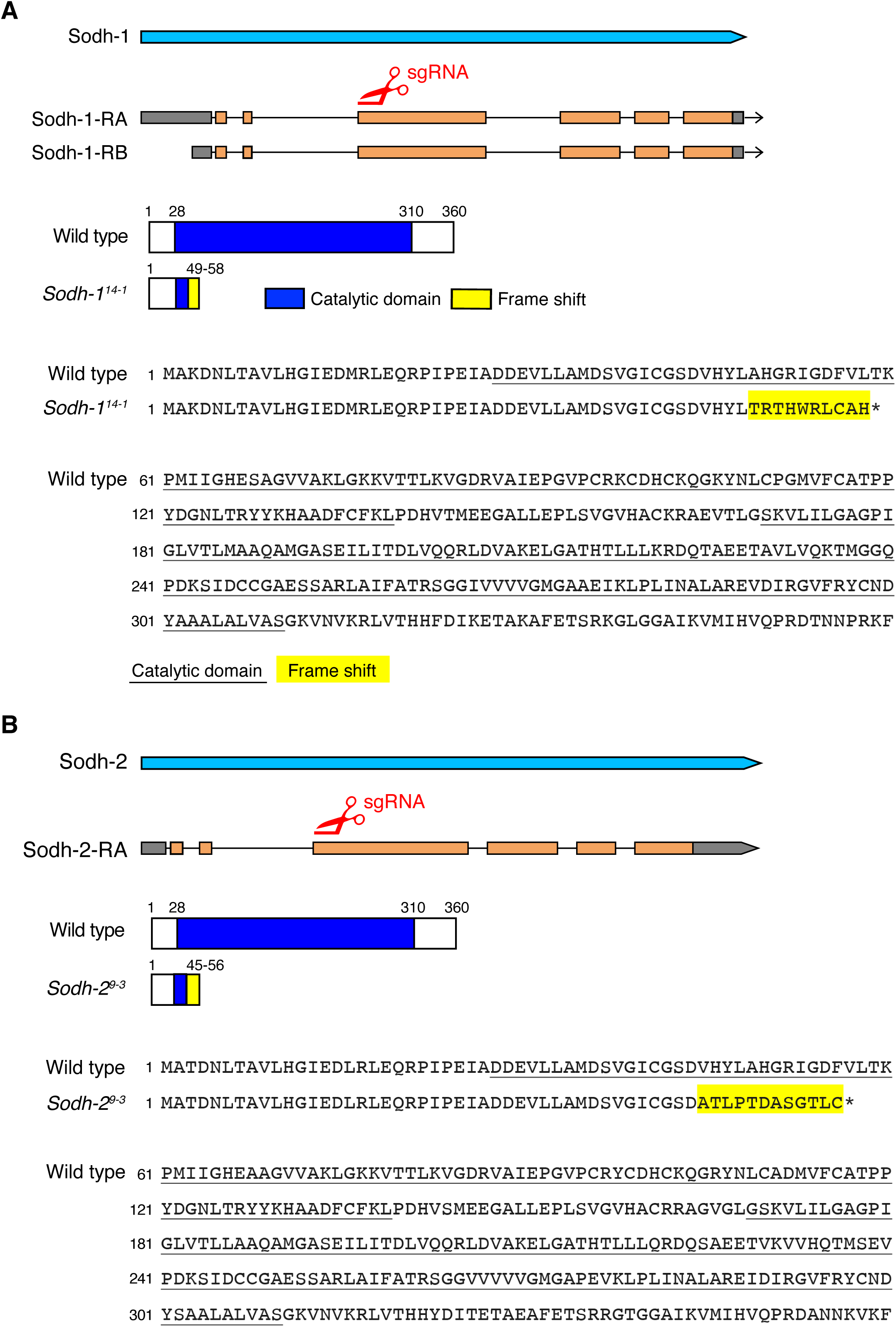
Generation of the *Sodh* mutant alleles. **A**, CRISPR-mediated mutagenesis of *Sodh-1*. A sgRNA was designed against the sequence in the exon common to both *Sodh-1* isoforms. The *Sodh-1*^*14-1*^ mutation caused a frame-shift (yellow) leading to premature termination of both isoforms of the Sodh-1 protein. The mutant proteins lack most of the catalytic domain (blue in schematic, underlined in sequence). **B**, CRISPR-mediated mutagenesis of *Sodh-2*. A sgRNA was designed against the sequence in the third exon of *Sodh-2*. The *Sodh-2*^*9-3*^ mutation caused a frame-shift (yellow) leading to premature termination of the Sodh-2 protein. The mutant proteins lack most of the catalytic domain (blue in schematic, underlined in sequence).

**Fig. S4.**
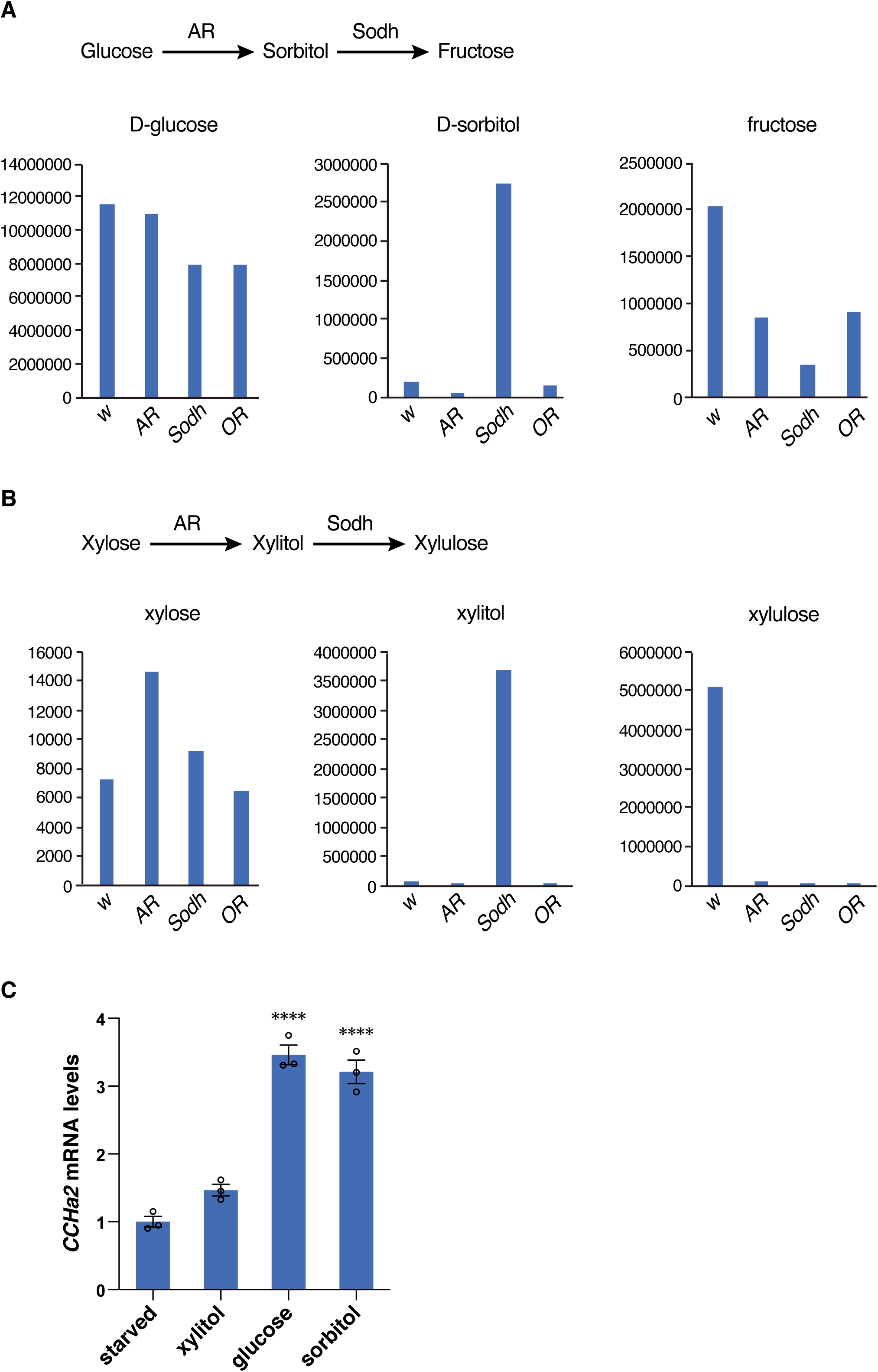
Metabolic phenotype of *AR* and *Sodh* mutants. **A**, The amount of glucose, sorbitol, and fructose contained in the hemolymph of *AR* and *Sodh* mutant third-instar larvae was measured using GS/MS. **B**, The amount of xylose, xylitol, and xylulose contained in the hemolymph of *AR* and *Sodh* mutant third-instar larvae was measured using GS/MS. *W* and *OR* were used as a control. **C**, Starved wild-type larvae were re-fed with 10% xylitol, glucose, or sorbitol for 6 hours. 10 larvae per batch, n=3 batches for all experiments. Histograms show mean ± SE. s\*\*\*\**P* < 0.0001.

**Fig. S5.**
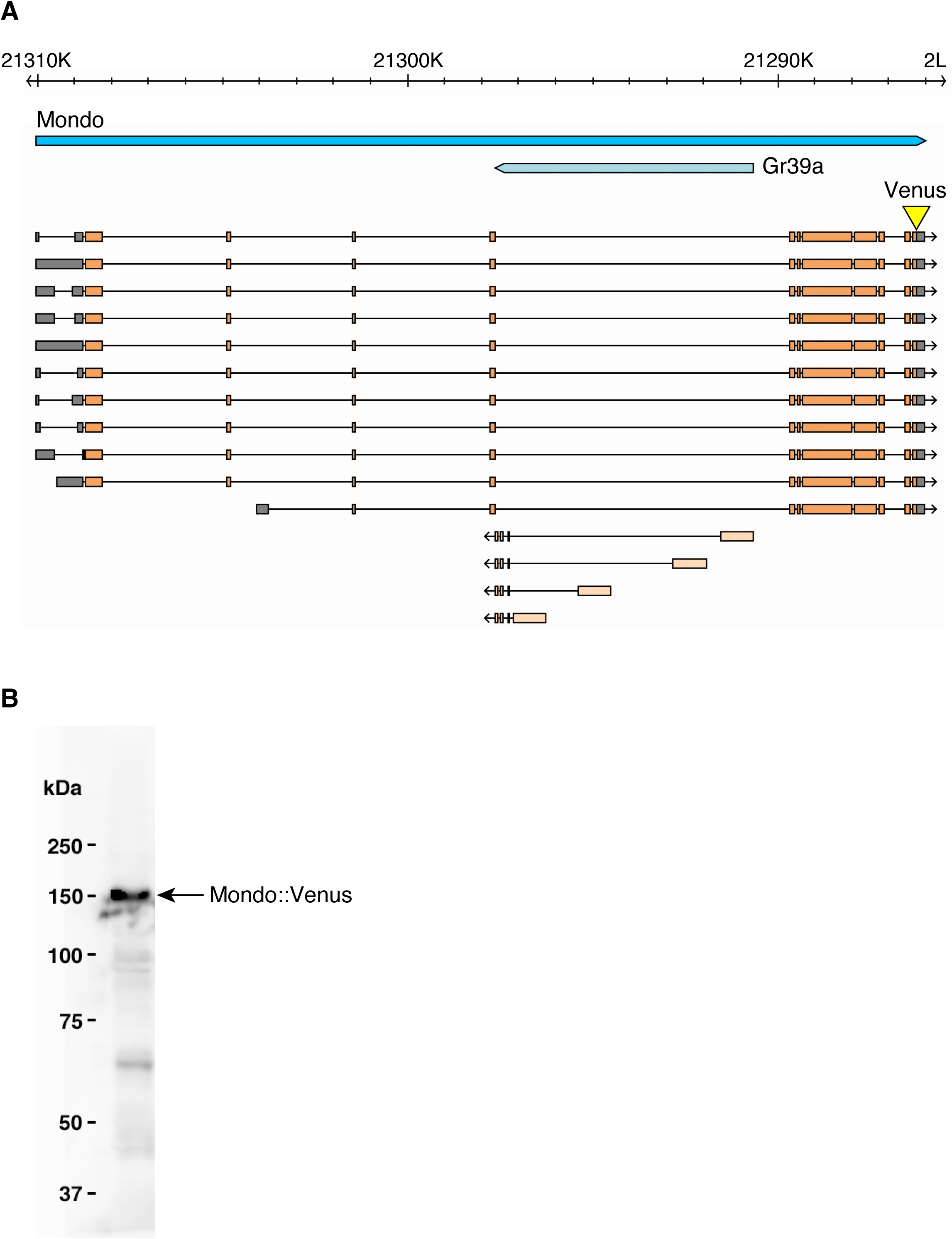
Knockin of the Venus fluorescent protein in the *Mondo* locus. **A**, Schematic drawing of the Mondo locus (adapted from FlyBase, http://flybase.org). The Venus fluorescent protein was knocked-in at the C-terminus of the Mondo coding region (yellow). **B**, Western blot using fat body extracts from the Mondo::Venus line. The Mondo::Venus fusion protein was detected using the anti-GFP polyclonal antibody.

**Fig. S6.**
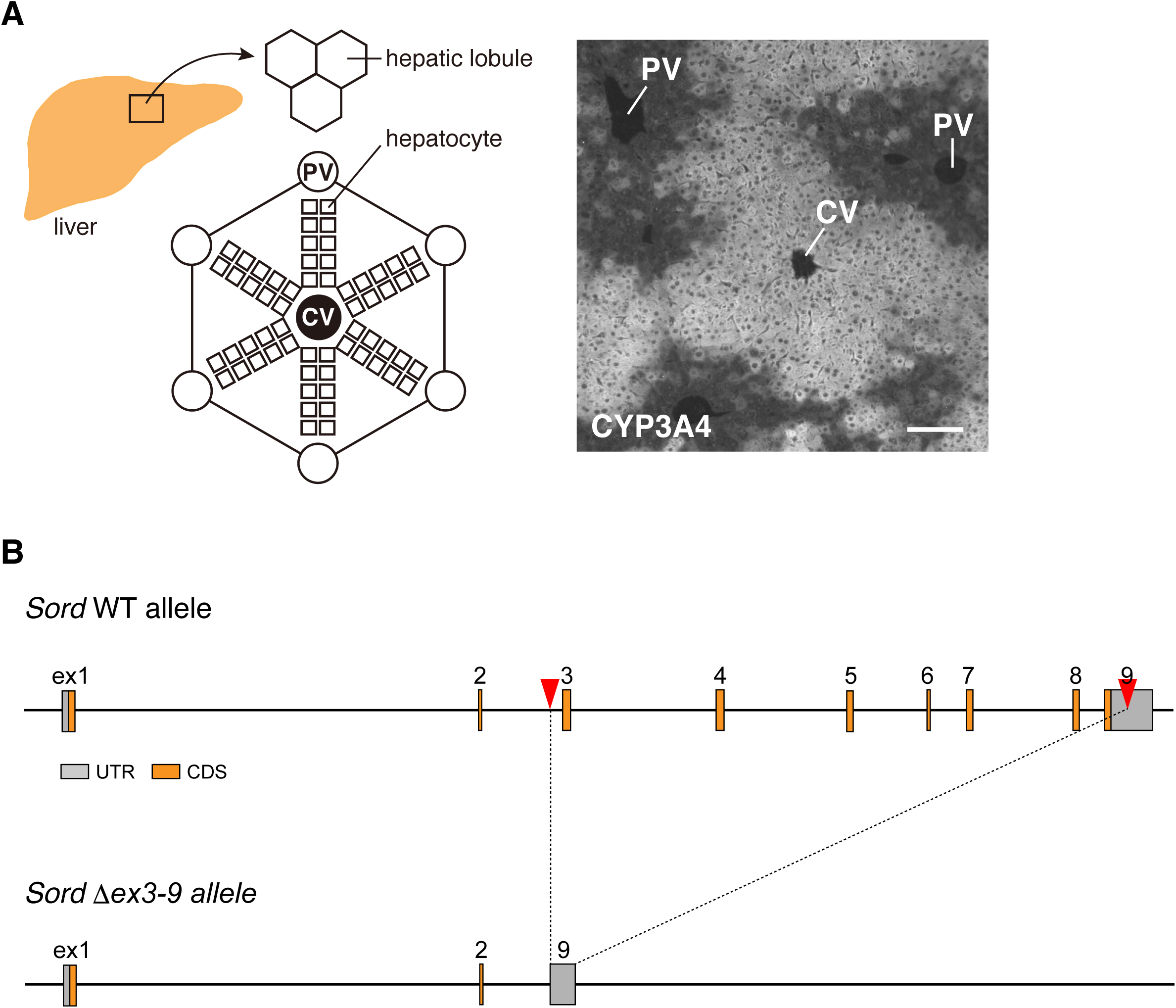
Analysis of ChREBP localization in mouse hepatocytes. **A**, Frozen liver sections were stained with anti-CYP3A4 antibody to label pericentral hepatocytes. Regions of interest were set on the CYP3A4-positive area for quantification of ChREBP signals in pericentral hepatocytes. Central vein (CV) and portal vein (PV) are indicated in the picture. **B**, CRISPR-mediated knockout of *Sord*. A crRNA was designed for the sequences in the intron 2 and the exon 9 of the *Sord* gene, resulting in the deletion from the exon 3 to the middle of the exon 9. Scale bar represents 100 µm.

**Fig. S7.**
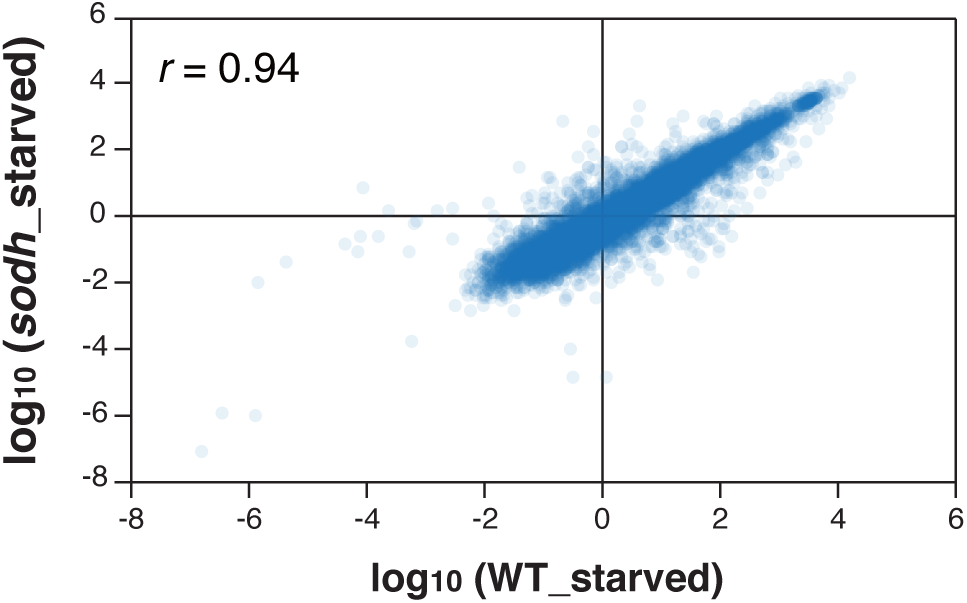
Transcriptomes of starved wild-type and *Sodh* mutant larvae. A comparison of the transcriptomes of starved wild-type and *Sodh* mutant larvae. 30 larvae per batch, n = 3 batches. Correlation coefficient (*r)* is indicated in the plot.

**Data S1. Sugar-responsive Mondo/Mlx-target genes (related to Fig. 3)**.

Previous study has reported sugar-dependent transcriptomes in wild-type and mutants of *max-like protein X* (*mlx*, also known as *bigmax*), the obligated partner of Mondo (Mattila et al., 2015). To identify Mondo-target genes, RNA-seq detasets reported in (GES70980) (Mattila et al., 2015) were analyzed using the FlyBase reference genome (Dmel Release 6.19). First, we selected genes whose expression levels were significantly changed between control and *mlx* mutant larvae under high sugar conditions. Of those, genes whose expression levels were significantly different between control and *mlx* mutants under low sugar conditions were removed. control_HSD = average expression levels of triplicated experiments of HSD-fed control larvae (FPKM), mlx1_HSD = average expression levels of triplicated experiments of HSD-fed *mlx* mutant larvae (FPKM), logFC = log2 fold-change, test_stat = test statistics, p_value = uncorrected p-value of the test statistics, q_value = adjusted p-value of the test statistics with Benjamin-Hochberg correction.

**Data S2. Differential expression test data of Mondo/Mlx-target genes in starved and glucose-fed wild-type larvae (related to Fig. 3)**.

WT_starved = average expression levels of triplicated experiments of starved wild-type larvae (FPKM), WT_glucose = average expression levels of triplicated experiments of glucose-fed wild-type larvae (FPKM), logFC = log2 fold-change, test_stat = test statistics, p_value = uncorrected p-value of the test statistics, q_value = adjusted p-value of the test statistics with Benjamin-Hochberg correction.

**Data S3. Differential expression test data of Mondo/Mlx-target genes in starved and sorbitol-fed wild-type larvae (related to Fig. 3)**.

WT_starved = average expression levels of triplicated experiments of starved wild-type larvae (FPKM), WT_sorbitol = average expression levels of triplicated experiments of sorbitol-fed wild-type larvae (FPKM), logFC = log2 fold-change, test_stat = test statistics, p_value = uncorrected p-value of the test statistics, q_value = adjusted p-value of the test statistics with Benjamin-Hochberg correction.

**Data S4. Differential expression test data of Mondo/Mlx-target genes in starved and sorbitol-fed *Sodh* mutant larvae (related to Fig. 3)**.

sodh_starved = average expression levels of triplicated experiments of starved *Sodh* mutant larvae (FPKM), sodh_sorbitol = average expression levels of triplicated experiments of sorbitol-fed *Sodh* mutant larvae (FPKM), logFC = log2 fold-change, test_stat = test statistics, p_value = uncorrected p-value of the test statistics, q_value = adjusted p-value of the test statistics with Benjamin-Hochberg correction.

**Data S5. Differential expression test data of Mondo/Mlx-target genes in starved and fructose-fed *Sodh* mutant larvae (related to Fig. 3)**.

sodh_starved = average expression levels of triplicated experiments of starved *Sodh* mutant larvae (FPKM), sodh_fructose = average expression levels of triplicated experiments of fructose-fed *Sodh* mutant larvae (FPKM), logFC = log2 fold-change, test_stat = test statistics, p_value = uncorrected p-value of the test statistics, q_value = adjusted p-value of the test statistics with Benjamin-Hochberg correction.

## Notes

### Competing Interest Statement

The authors have declared no competing interest.

